# C3G (RapGEF1) localizes to the mother centriole and regulates centriole division and primary cilia dynamics

**DOI:** 10.1101/575225

**Authors:** Sanjeev Chavan Nayak, Vegesna Radha

**Affiliations:** Centre for Cellular and Molecular Biology, Uppal Road, Hyderabad – 500 007, INDIA

**Keywords:** C3G, RapGEF1, Centrosome, Cenexin, Primary cilia, Centriole

## Abstract

C3G (RapGEF1), a negative regulator of β-catenin, plays a role in cell differentiation and is essential for early embryonic development in mice. In this study, we identify C3G as a centrosomal protein that regulates centriole division and primary cilia dynamics. C3G is present at the centrosome in interphase as well as mitotic cells, but is absent at the centrioles in differentiated myotubes. It interacts with, and co-localizes with cenexin in the mother centriole. Stable clone of cells depleted of C3G by CRISPR/Cas9 showed reduction in cenexin protein, and presence of supernumerary centrioles. Over-expression of C3G resulted in inhibition of centrosome division in normal and hydroxyurea treated cells. Proportion of ciliated cells is higher, and cilia length longer in C3G knockout cells. C3G inhibits cilia formation and length dependent on its catalytic activity. Unlike wild type cells, C3G depleted cells inefficiently retracted their cilia upon stimulation to reenter the cell cycle, and proliferated slowly, arresting in G1. We conclude that C3G inhibits centriole division and maintains ciliary homeostasis, properties that may be important for its role in embryonic development.

**Summary statement:** We identify C3G as a centrosomal protein and regulator of centriole number, primary cilia length and resorption. These properties are important for its role in embryogenesis, and suggest that mutations in C3G could cause ciliopathies.

## Introduction

Temporal regulation of cell fate in multicellular organisms during development and in adult tissues is dependent on signaling molecules that respond to distinct environmental cues which are translated to altered gene expression and cytoskeletal remodeling. Guanine nucleotide Exchange Factors (GEFs) that activate small GTPases are important mediators in signaling pathways and act as hubs to receive signals and relay them to specific effector functions (Overbeck et al., 1995; Quilliam et al., 2002; Rossman et al., 2005). The ubiquitously expressed protein, C3G (RapGEF1), regulates cell proliferation and differentiation, and is highly conserved across vertebrate phyla (Radha, 2011). It is a 140 kDa protein with a catalytic domain at the C terminus and a central region containing multiple polyproline tracts called Crk binding region (CBR). It activates GTPases of Ras & Rho family and interacts with several proteins like Crk, SFKs, c-Abl, TCPTP & β-catenin (Dayma et al., 2012; Gotoh et al., 1995; Knudsen et al., 1994; Mitra et al., 2011; Mochizuki et al., 2000; Radha et al., 2007; Shivakrupa et al., 2003). It can participate in signaling as a GEF as well as an adaptor molecule. Its GEF activity is regulated by protein interaction, membrane localization and phosphorylation (Ichiba et al., 1999; Ichiba et al., 1997; Sakkab et al., 2000). C3G phosphorylation at Y504 in human cells enhances its catalytic activity at distinct subcellular compartments (Mitra et al., 2011; Mitra and Radha, 2010; Radha et al., 2004).

C3G is involved in cytoskeletal remodeling and is required for neuronal and myogenic differentiation (Mitra et al., 2011; Radha et al., 2011; Sasi Kumar et al., 2015). Its loss causes embryonic lethality in mice indicating that early embryonic development is dependent on C3G functions (Ohba et al., 2001). Introduction of a hypomorphic allele prolongs survival but the embryos show defects in development of brain and other organ systems (Ohba et al., 2001; Voss et al., 2003). We have earlier shown that C3G interacts with, and regulates stability and activity of β-catenin (Dayma et al., 2012). In addition, C3G expression is regulated by β-catenin activity. Endogenous as well as exogenously expressed C3G shows prominent cytoplasmic localization, but it undergoes regulated nuclear translocation in response to physiological stimuli (Hogan et al., 2004; Radha et al., 2004; Shakyawar et al., 2017). Nuclear functions like chromatin remodeling and splicing are regulated by C3G (Shakyawar et al., 2017; Shakyawar et al., 2018). In a cell and tissue type dependent manner, C3G functions to either increase or decrease cell proliferation, and altered C3G levels are associated with human tumors and other disorders (Che et al., 2015; Guerrero et al., 1998; Gutiérrez-Berzal et al., 2006; Hirata et al., 2004; Ishimaru et al., 1999; Okino et al., 2006; Radha et al., 2011; Samuelsson et al., 2011; Sequera et al., 2018; Voss et al., 2006).

The centrosome with its two centrioles plays a pivotal role in cell division, and maintaining its numbers is important for proper chromosome segregation (Bettencourt-Dias and Glover, 2007). The two centrioles are asymmetric, and the mother centriole has appendages which aid in membrane anchoring and growth of primary cilia, a non motile microtubule based structure (Nigg and Stearns, 2011; Pelletier and Yamashita, 2012; Yadav et al., 2016). In cells that exit the cell cycle, the mother centriole matures into the basal body from which the primary cilium protrudes out of the cell membrane. It is retracted when cells reenter the cell cycle, and is therefore dynamically regulated (Avidor-Reiss and Gopalakrishnan, 2013; Goetz and Anderson, 2010; Kim and Dynlacht, 2013; Pan et al., 2004; Sanchez and Dynlacht, 2016). It serves as a signaling hub and plays an important role in cell fate decision during development (Eggenschwiler and Anderson, 2007; Goetz and Anderson, 2010; Hilgendorf et al., 2016). Defects in the structure and functions of primary cilium are associated with developmental defects in humans called as ciliopathies (Gerdes et al., 2009; Habbig and Liebau, 2015; Nigg and Raff, 2009). A large number of proteins make up the centrosome and mutations in centrosomal proteins, or their deregulation, cause a variety of human disorders. While proteins in the centrosome-cilium interface have been mapped, how they function to regulate appropriate centrosomal division, protrusion and resorption of primary cilia is not fully understood (Hsu et al., 2017; Gabriel et al., 2016; Gupta et al., 2015).

Centrosomal proteins tend to be larger than generic human proteins, since their genes contain on average more exons (20.3 versus 14.6). They are rich in predicted disordered regions, which cover 57% of their length, compared to 39% in the general human proteome (Dos Santos et al., 2013). The gene encoding C3G shares these properties in having 24 exons, and C3G protein has a highly disordered predicted structure. As a component of multiple signaling pathways important for cell fate decisions like proliferation and differentiation, we examined if C3G localizes to the centrosome and basal body. We show that C3G localizes to the sub distal appendages of the mother centriole in interphase cells and functions to regulate centriole duplication. CRISPR/CAS9 mediated knockout of C3G caused an increase in number of cells with cilia, as well as cilia length. Cells lacking C3G are unable to resorb their cilium upon stimulation to re-enter the cell cycle, suggesting that C3G is required for maintenance of cilia length and timely disassembly.

## Results

### C3G localizes to the centrosome

Cellular, as well as exogenously expressed C3G predominantly localizes to the cytoplasm of exponentially growing cells (Hogan et al., 2004; Radha et al., 2004). In response to nerve growth factor (NGF) treatment of IMR-32 neuroblastoma cells, C3G is phosphorylated at Y504 and localizes to the core of the Golgi (Mitra et al., 2011; Radha et al., 2008). In interphase cells, the Golgi and centrosome are located adjacent to each other, close to the nucleus (Sutterlin and Colanzi, 2010) prompting us to examine the localization of endogenous C3G to the centrosome by confocal microscopy. Towards this end, we used a commercial antibody raised against sequences in the N-terminus of C3G which specifically recognizes a 140 kDa polypeptide specific to C3G in a variety of cell types (Fig.S1A) (Begum et al., 2018; Guerrero et al., 2004; Radha et al., 2007; Shakyawar et al., 2017). Z stacks of methanol fixed C2C12 cells showed C3G staining as a single sharp dot of about 0.5 μm diameter positioned juxta nuclear in each cell (Fig.S1B). This staining was lost in cells when C3G expression was knocked down using ShRNA (Fig.S1C & S1D). γ-tubulin antibody marks the centrioles within the centrosome and is seen as two closely spaced spots in interphase cells (Vorobjev et al., 2000). The juxta nuclear dot like pattern of staining seemed consistent with the presence of C3G in the centrosome, and its co-localization with γ-tubulin confirmed its association with the centriole in C2C12 and ARPE-19 cells (Fig.1A). To further confirm localization to the centriole, we examined staining in a clone of cells lacking C3G due to CRISPR/CAS9 mediated knockout (Shakyawar et al., 2017). These cells did not show a signal colocalizing with γ-tubulin (Fig.1B). Reduction in C3G levels was validated by western blotting (Fig.1C). An alternate C3G antibody (rGRF2) also confirmed localization of C3G at the centrosome (Fig.S1E). In addition, co-staining of cells for C3G and PCM1, a protein in the pericentriolar material of the centrosome (Ou et al., 2004) showed localization of C3G at the core of the centrosome (Fig.S1F). Endogenous C3G in C2C12 cells colocalized with GFP-centrin, a marker of centrioles (Fig.S1G). Overexpressed C3G-GFP also co-localized with γ-tubulin at the centrosome, in addition to its prominent presence throughout the cytoplasm (Fig.S1H). These results demonstrated that C3G specifically localizes to the centrosome.

**Figure 1.**
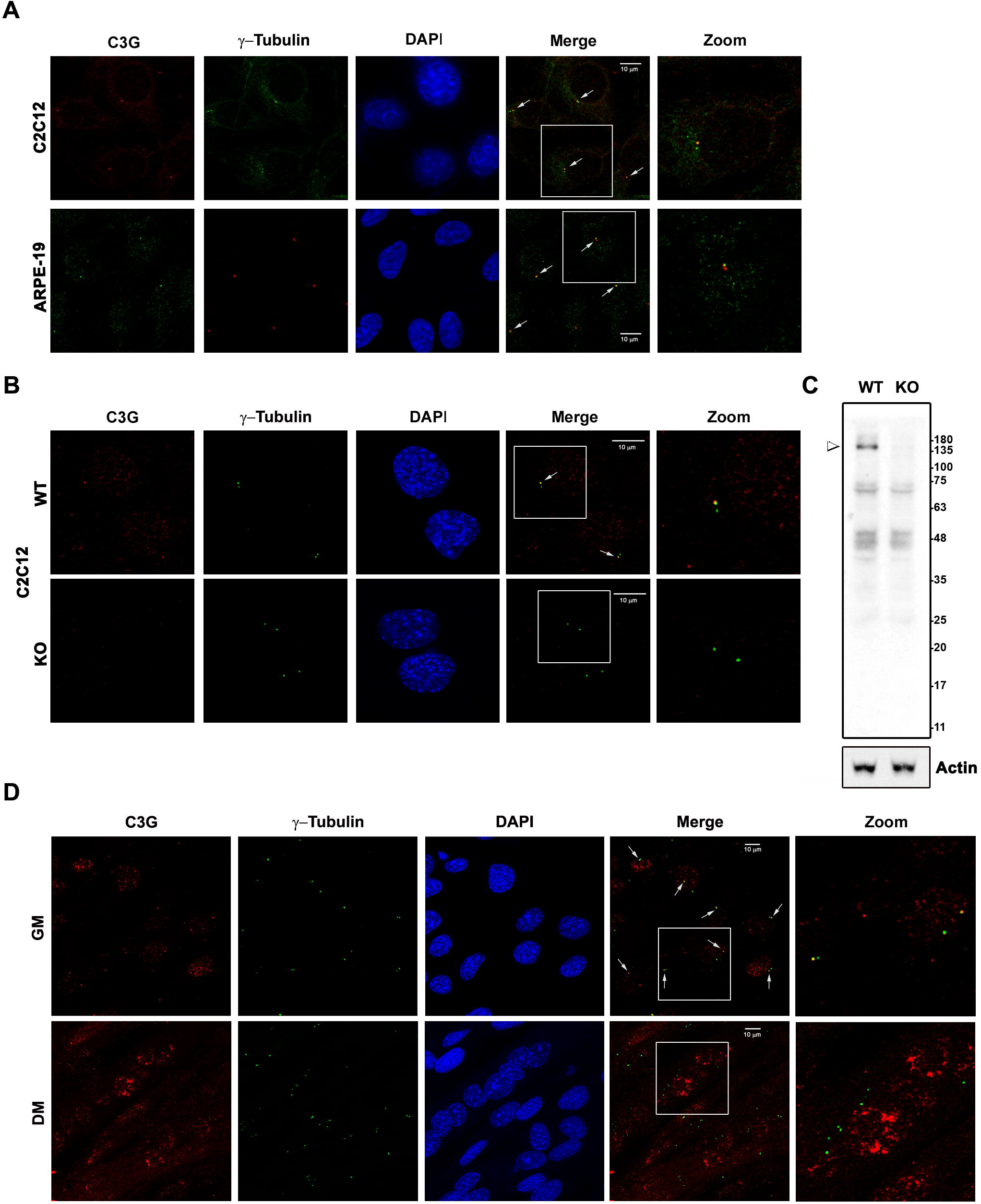
C3G localizes to the centrosome. A. Exponentially growing C2C12 and ARPE-19 cells were examined for endogenous C3G and γ-tubulin localization by indirect immunofluorescence. Panels show a single Z-section across the centrosome captured using a 63X objective. B. C2C12 WT and C3G KO clones were dual stained for C3G and γ-tubulin. C. Western blot showing C3G levels in WT and KO cells. Actin was used as loading control. Arrowhead points to the 140 kDa C3G polypeptide. D. C2C12 cells grown in growth medium or in differentiation medium for 96 hours were stained for C3G and γ-tubulin. In A, B & D, the areas zoomed into are indicated, and arrows point to C3G colocalizing with γ-tubulin at the centrosome.

Physiological cell fusion during skeletal muscle differentiation results in multinucleated cells with several centrioles (Bugnard et al., 2005). While undifferentiated myocytes have a single nucleus and a pair of centrioles, myotubes possess multiple centrioles which congregate close to the nuclei. C3G localization to the centrioles was examined in myotubes formed, in culture, upon differentiation of C2C12 cells. Unlike in undifferentiated cells, C3G did not colocalize with γ-tubulin at centrioles in myotubes (Fig.1D). As shown earlier (Shakyawar et al., 2017), C3G localized prominently to the nuclei of myotubes. C3G localization to the centrosome may be dynamic and dependent on the physiological state of the cell.

Many proteins are known to show dynamic localization to the centrosome (Hames et al., 2005; Lui et al., 2016). Since C3G is primarily present in the cytoplasm, we examined if its presence at the centrosome is dependent on intact cytoskeletal structures in the cell. Cells were treated with cytochalasin D (which disrupts actin microfilaments) or nocodazole (which disrupts microtubules) and examined for C3G localization to centrosome. We observed that centrosomal localization of C3G was disrupted in cells treated with either of these drugs (Fig.S2A). Nocodazole treated cells also showed weak staining for γ-tubulin, as described earlier (Vorobjev et al., 2000). The efficacy of the drugs used is shown in (Fig.S2B & S2C). Nocodazole disrupted α-tubulin structures, and cytochalasin disrupted F-actin.

### Localization of C3G and phosphorylated C3G (active) to mitotic centrosomes

Localization of proteins to the centrosome may change as cells enter and exit mitosis (Hames et al., 2005; Lu et al., 2009; Lui et al., 2016). C2C12 cells stained for C3G and γ-tubulin showed distinct staining for C3G at the centrosomes at various phases of mitosis (Fig.2). Interestingly, interphase cells showed association of C3G with only one of the centrioles. In early prophase, where the newly divided centrosomes have not moved to the poles, C3G was associated with one of the centrosomes, but was present in both centrosomes in all other mitotic phases. C3G staining was more prominent in the centrosomes of mitotic cells, compared to that seen in interphase cells. Enhanced centrosomal C3G staining seen in mitotic phases reverted back to that seen in interphase cells during cytokinesis. Co-localization was also observed with GFP-Centrin, during mitosis (Fig.S1G).

**Figure 2.**
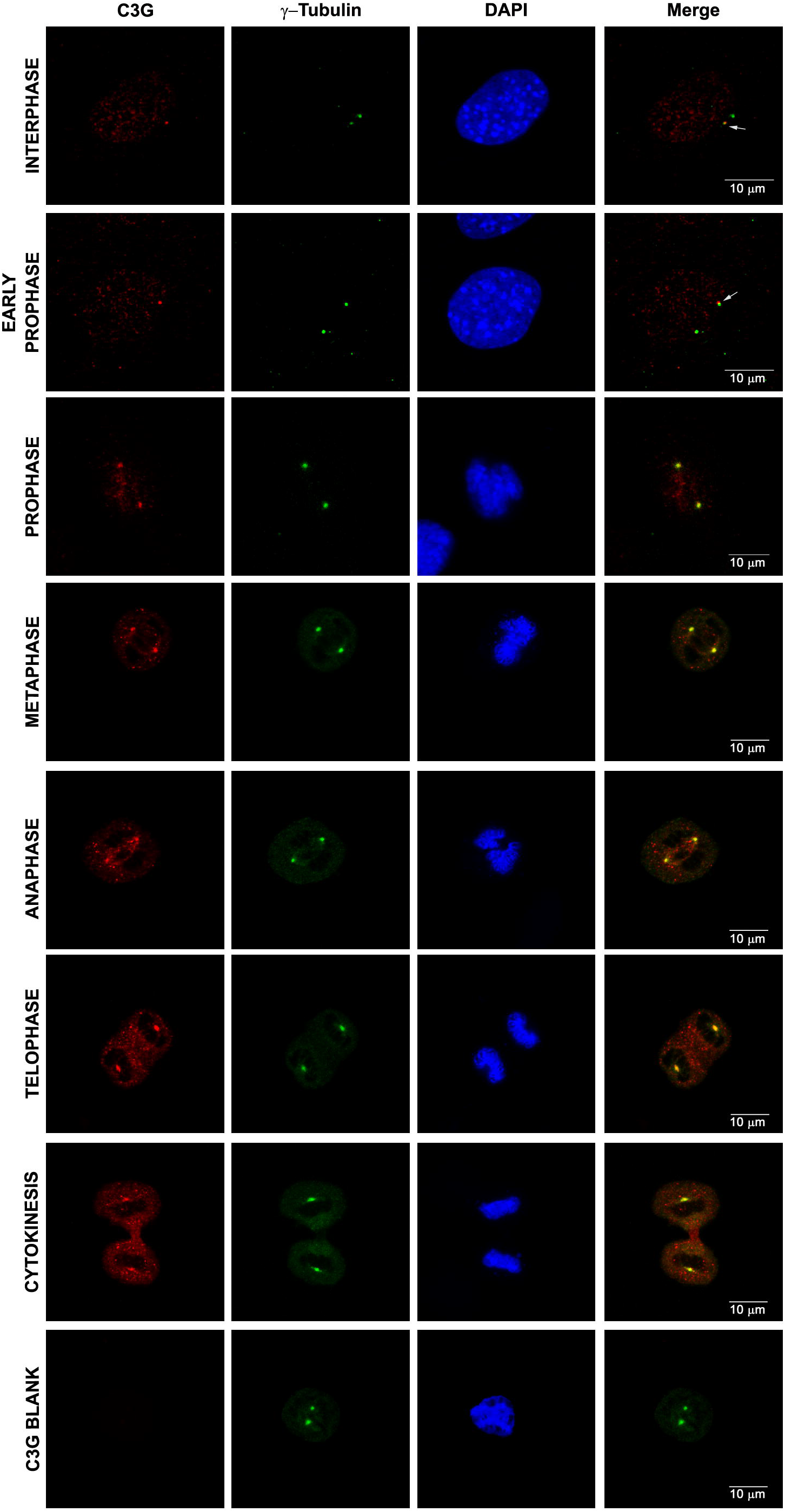
C3G localizes to the centrosome during mitosis. Exponentially growing C2C12 cells were dual-stained by indirect immunofluorescence for C3G and γ-tubulin. Cells in various phases of mitosis were identified based on morphology and DAPI staining, and images captured on Leica SP8 confocal microscope. Panels show merged images of optical sections of z-plane step size of 0.35 μm, captured across the cell diameter using a 63X objective. C3G blank indicates cells processed similarly without the addition of primary antibody against C3G. Arrows point to C3G staining associated with one of the centrioles in interphase and early prophase.

Phosphorylation (Y504 in human; Y514 in mouse), increases the GEF activity of C3G towards Rap GTPase (Ichiba et al., 1999). Upon inhibition of cellular tyrosine phosphatases by pervanadate (PV), expression of Src family kinases, or c-Abl; phosphorylated or active C3G is seen in distinct sub cellular regions (Mitra et al., 2011; Radha et al., 2008; Radha et al., 2004). pY514 C3G in C2C12 cells co-localized with γ-tubulin in mitotic cells, with barely detectable signals seen in interphase cells (Fig.S3A). These results indicate that catalytically active C3G is present particularly at the centrosome in cells undergoing mitosis, although C3G is seen at interphase and mitotic centrosomes. Co-localization of pC3G with γ-tubulin was also observed in PV treated IMR-32 cells, though the signal was diffused and seen in the region surrounding the centrioles (Fig.S3B). Increase in pC3G upon PV treatment was confirmed by western blotting (Fig.S3C). This pattern of staining was similar to that seen in NGF or forskolin treated IMR-32 cells (Mitra et al., 2011; Radha et al., 2004). These results indicated that activation of C3G at the centrosome in interphase cells may be transient, occurring in response to distinct signals, and can be detected only upon inhibition of cellular tyrosine phosphatases.

### C3G localizes to the mother centriole

The presence of C3G staining with one of the interphase centrioles suggested its differential localization to the two centrioles. We examined its co-localization with γ-tubulin at higher magnification and confirmed that C3G is associated with only one of the centrioles in C2C12 as well as ARPE-19 cells (Fig.3A & 3B). Intensity scanning of an ROI across the two centrioles showed C3G staining with only one of the γ-tubulin intensity peaks in both cell types (Fig.S4A & S4B). C3G staining was seen at one end of γ-tubulin suggesting a pattern shown by appendage proteins of the mother centriole.

Localization of C3G to the mother centriole was examined by transiently expressing GFP-Cenexin, a marker for centriolar sub distal appendages (Lange and Gull, 1995; Nakagawa et al., 2001). Endogenous C3G co-localized with GFP-Cenexin (Fig.3C). Over-expression of GFP-Cenexin is known to form aggregates (Steere et al., 2012), which were observed in some of the transfected cells. Interestingly, we found that endogenous C3G also co-localizes with these structures, which appear as rings in confocal sections. The intensity of C3G staining in these structures was far more than at the centrioles, suggesting that significant amount of cellular C3G co-localizes with cenexin when it forms aggregates, due to over-expression. An alternate antibody to C3G (rGRF2) showed colocalization with GFP in cells transfected with GFP-Cenexin, and with endogenous cenexin (Fig.S4C & S4D). Intensity scanning of an ROI across the cenexin signal showed that the signal intensity of C3G matched that of cenexin (Fig.S4E). These results showed that C3G is localized to the sub-distal appendages of the mother centriole and suggested its possible interaction with cenexin. To examine this, we expressed C3G-Flag & GFP-Cenexin in HEK293T cells and carried out immunoprecipitation of C3G. As shown in (Fig.3D), cenexin was present in the complex pulled down with C3G antibodies, but not in complex with normal IgG. Interaction was further confirmed by pulling down cenexin from cells transfected with GFP-Cenexin and C3G-Flag vectors, using GFP-Trap beads. C3G was present in complex with GFP-Cenexin but not GFP (Fig.3E). These results indicated that C3G and cenexin are part of a common complex in cells.

**Figure 3.**
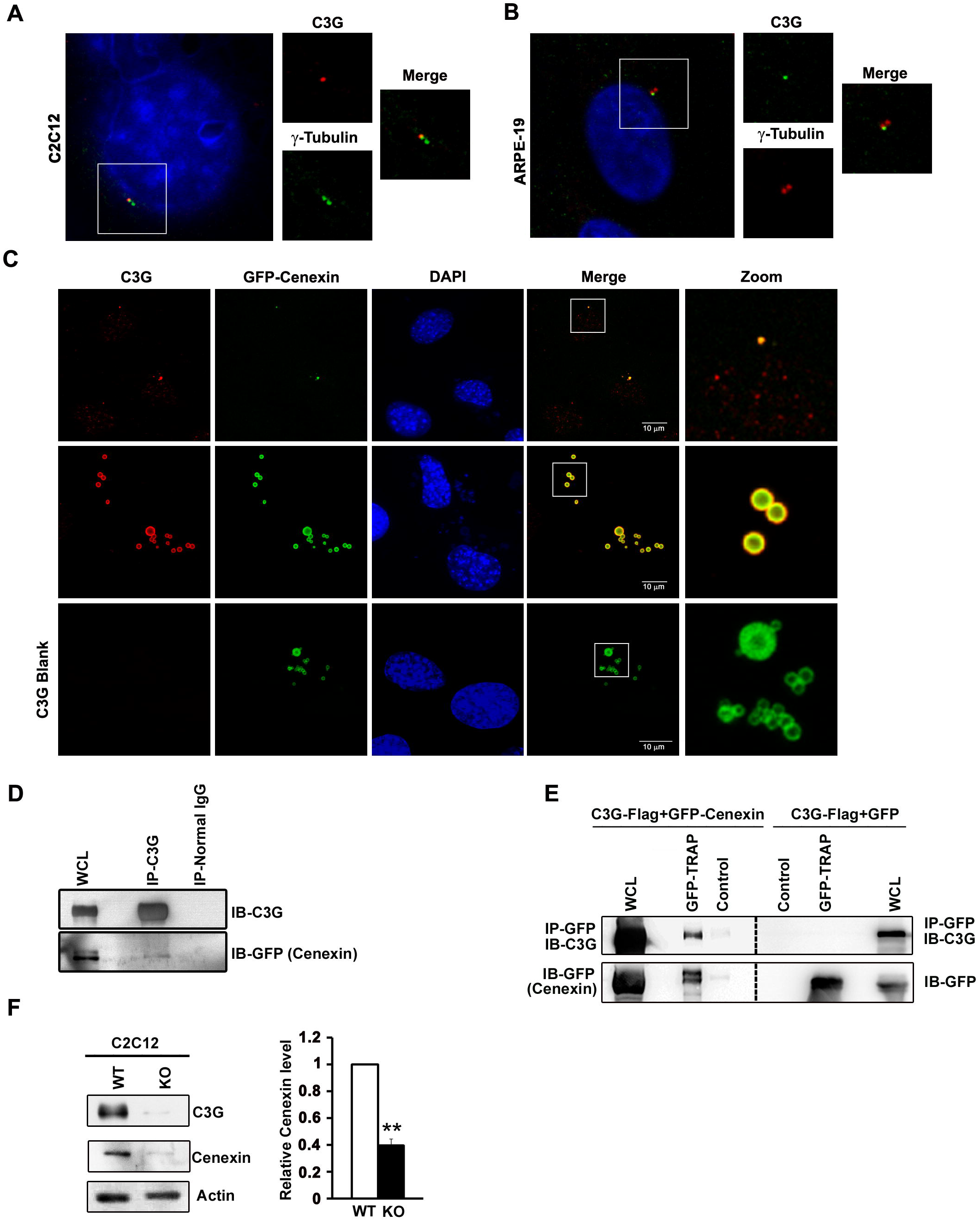
C3G Localizes to the mother centriole. A,B. High magnification images showing merged optical sections across the centrosome dually-labelled with antibodies against C3G and γ-tubulin, showing C3G co-staining with only one of the γ-tubulin spots in C2C12 and ARPE-19 cells. C.C2C12 cells were transfected with GFP-cenexin, a sub-distal appendage protein of the mother centriole, and stained with C3G antibody. Upper panel shows cells with moderate GFP-Cenexin expression localizing to a single centriole. Middle panel shows cells with high expression, where they form aggregates. C3G blank indicates cells processed similarly, but without addition of C3G antibody. D.C3G interacts with cenexin. HEK293T cell lysate coexpressing C3G-Flag and GFP-Cenexin was subjected to immunoprecipitation using antibody against C3G. Normal mouse IgG was used as control. Western blotting was carried out using GFP antibody to detect co-precipitation of cenexin. E. HEK293T cell lysates co-expressing C3G-Flag and GFP-Cenexin, or C3G-Flag and GFP (control) were used to immunoprecipitate GFP-Cenexin using agarose-conjugated GFP antibody (GFP Trap) or control beads. Western blotting was carried out to detect co-immunoprecipitation of C3G. F. Cellular cenexin levels are reduced in C3G KO cells. Western blot showing endogenous cenexin in lysates of C2C12 WT and KO cells. Actin was used as a loading control. Bar diagram shows cenexin protein levels quantified from three independent experiments. ** p <0.01.

Cenexin has been shown to be required for ciliogenesis and regulation of centrosome division (Yang et al., 2018; Ishikawa et al., 2005). Its transcription is up regulated in quiescent cells (Pletz et al., 2013). Anchoring of proteins at the appendages is dependent on molecular interactions with cenexin (Hehnly et al., 2013; Ishikawa and Marshall, 2011; Soung et al., 2009). Since C3G colocalizes with cenexin, we examined cenexin levels in cells lacking C3G. Cellular cenexin protein levels are significantly reduced in C2C12 clones lacking C3G expression (Fig.3F). This result was also confirmed in human cells using multiple clones of C3G knockout (KO) MDA-MB-231 cells, generated using CRISPR/CAS9. Compared to wild type (WT) cells, C3G KO clones showed reduced cenexin levels that correlated well with reduction in C3G levels (Fig.S7A). Over-expression of C3G caused a small increase in cellular cenexin levels in C2C12 as well as MDA-MB-231 cells (Fig.S7C), suggesting that C3G regulates cellular cenexin levels.

### C3G regulates centriole division

Proteins that localize to the mother centriole play a role in centrosome division (Boutros and Ducommun, 2008), and optimal cenexin levels are required to regulate centrosome division (Yang et al., 2018). We therefore examined if loss of C3G has an effect on centriole number. WT & C3G KO C2C12 cells were stained for γ-tubulin and examined for the number of centrioles in interphase cells. A large number C3G KO cells showed the presence of supernumerary centrioles (Fig.4A). Quantitation showed that the percentage of interphase cells with more than two γ-tubulin spots was higher in KO cells when compared to WT cells (Fig.4A, bar diagram).

**Figure 4.**
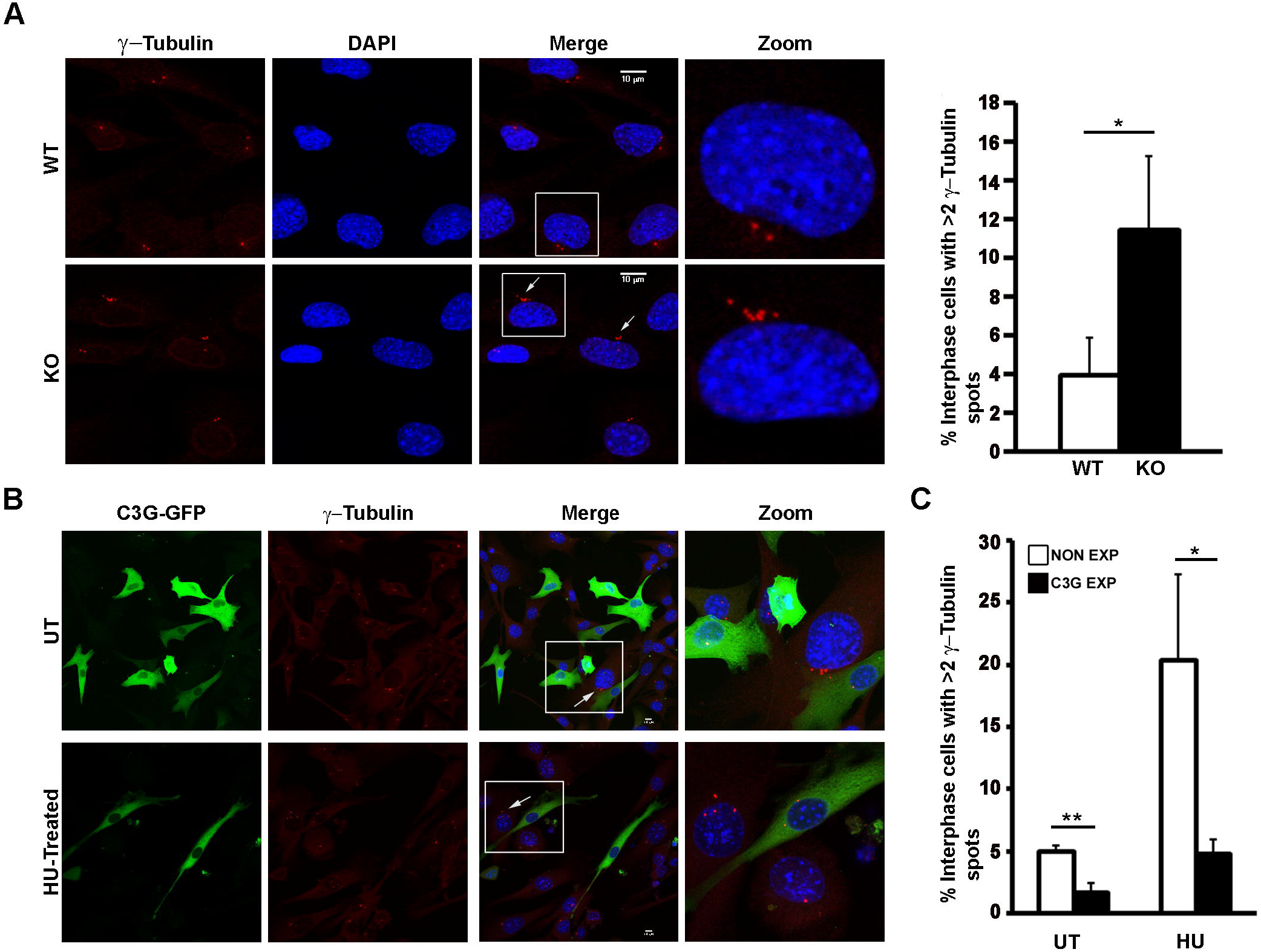
C3G regulates centriole division. A. C2C12 WT and C3G KO cells were grown on coverslips and stained with anti γ-tubulin antibody to examine centriole number. Panels show the presence of supernumerary centrioles in KO cells. Bar graph shows the percentage of interphase cells with >2 γ-tubulin spots, from three independent experiments carried out on duplicate cover slips. *p<0.05. B. C3G over-expression inhibits centrosome division. C2C12 cells transfected with C3G-GFP were left untreated (UT), or treated with hydroxyurea (HU) and stained for C3G anti γ-tubulin. Arrows point to supernumerary centrioles. C. Bar graph shows the percentage of interphase cells with >2 γ-tubulin positive spots in expressing and nonexpressing cells. Data are expressed as means ± standard deviations of at least 50 expressing and 100 non-expressing cells in each duplicate cover slip from 3 independent experiments. *p<0.05, **p<0.01.

If loss of C3G causes abnormal centriole division, we presumed that increase in its levels may also alter centrosome division. Treatment with hydroxyurea (HU) inhibits nuclear division, but not centriolar division, resulting in cells with multiple centrioles (Balczon et al., 1995). Transiently transfected C3G-GFP cells were fixed and stained for γ-tubulin to visualize centrioles in the presence or absence of HU. Proportion of interphase cells with more than two centrioles was quantitated among C3G expressing and non expressing cells. C3G over expression was inhibitory to centrosome division, both in normally growing cells and in HU treated cells (Fig.4B). Quantitation indicated that fewer C3G expressing cells showed supernumerary centrioles compared to non-expressing cells (Fig.4C). Similar effect of C3G over-expression on centriole number was seen in IMR-32 cells (Fig.S5). Therefore, optimal levels of C3G appear to be required for maintaining normal centriole number.

### C3G localizes to the basal body and regulates primary cilia

The mother centriole in cells that exit the cell cycle forms the basal body which gives rise to the primary cilium, which can be detected by acetylated tubulin (Ac-tub) staining. C2C12 cells which put forth a primary cilium upon serum starvation, showed the presence of C3G at the basal body (Fig. 5 A). Localization of C3G at the basal body was also seen in serum starved ARPE-19 cells (data not shown). C3G KO C2C12 clones did not show any staining at the basal body, though primary cilia were present in these cells (Fig.5B). Reduced C3G levels in the KO clone used is shown in the blot (Fig.5C). Staining for Ac-Tub was intact in C3G KO cells, but we observed a difference in the morphology of primary cilia, which appeared longer (Fig.5B). We quantitated the number of cells with primary cilia and also its length in C3G KO clones and compared them with WT cells. This was carried out under condition of exponential growth, where only a small fraction of C2C12 cells show cilia, and also under condition of serum starvation for 24 hrs when cells put forth cilia. Under these conditions, cells exit the cell cycle, but do not fuse to form myotubes. Fig.5B shows representative images of WT & KO cells; quantitation is shown in Fig.5D. C3G KO clone shows a significantly higher proportion of cells with primary cilium in exponentially growing as well as serum starved conditions, compared to the WT. Also, cilia were significantly longer in KO cells (Fig.5E).

**Figure 5.**
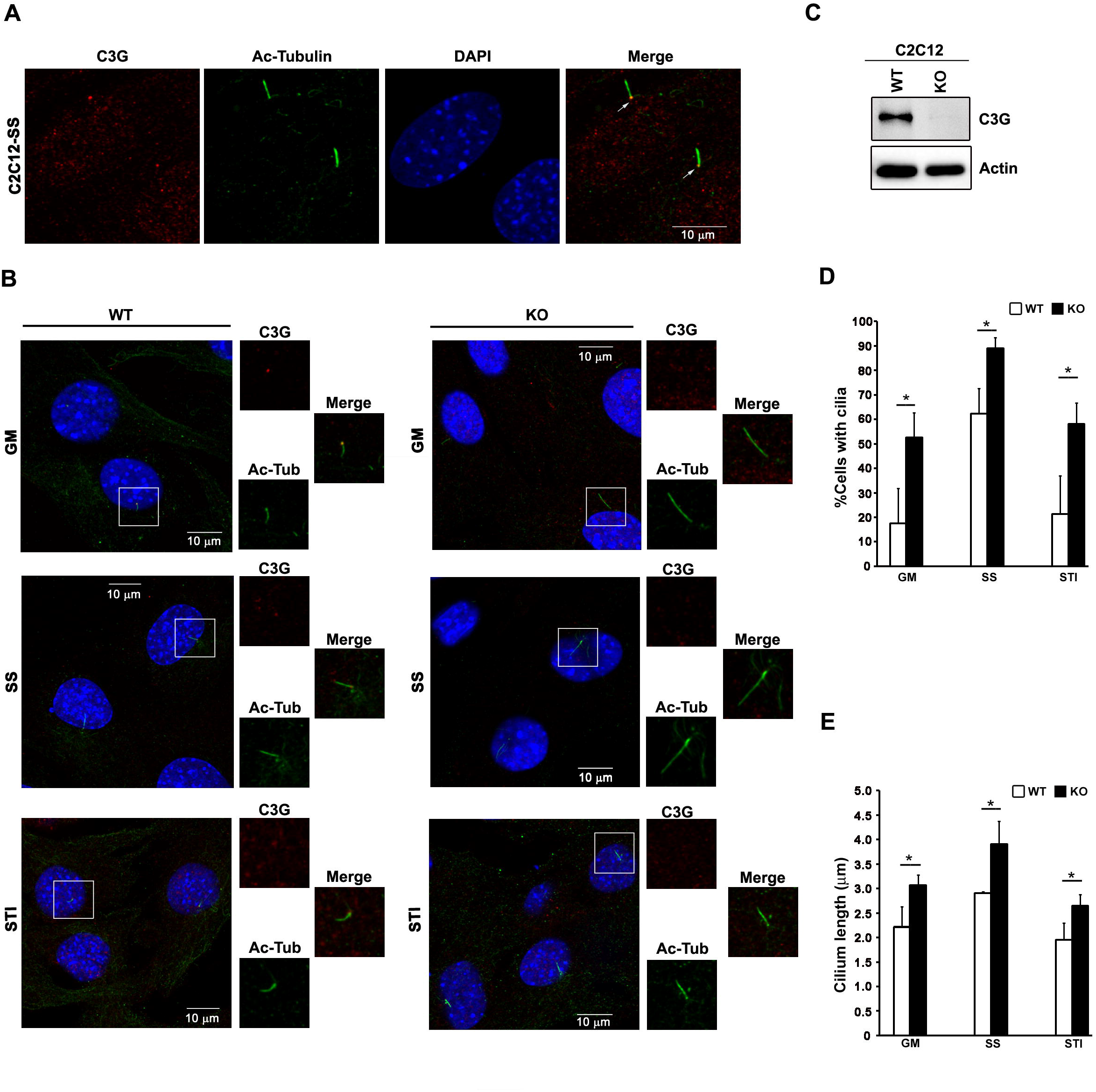
C3G localizes to the basal body and regulates primary cilia. A. C2C12 cells were serum-starved, and indirect immunofluorescence carried out to visualize C3G and Ac-tubulin, a marker of primary cilia. Arrows point to C3G presence at the basal body. B. WT and KO C2C12 cells were cultured in growth media (GM) or serum-starved (SS), or re-stimulated after serum starvation (STI). Cells were dual-stained for C3G and Ac-tubulin, and examined for presence of primary cilia. Panels show fields of cells with primary cilia. C. Western blot showing C3G protein level in WT and C3G-KO C2C12 cells. Actin was used as a loading control. D. Bar graph shows percentage of WT and KO C2C12 cells having cilia quantitated from three independent experiments carried out as described in (B). *p <0.05. E. Bar graph shows the average cilium length in WT and KO C2C12 cells cultured under conditions of growth, serum starvation and re-stimulation conditions from three independent experiments. *p<0.05.

### C3G knockout cells are compromised for cilia resorption as cells re-enter the cell cycle

Primary cilia are dynamic structures that are formed and retracted specifically in a cell cycle dependent manner. Loss of C3G leading to enhanced cilia formation and longer cilia could be because C3G functions as a negative regulator of ciliogenesis by restricting cilia length, and aiding in their resorption. Primary cilia can be induced to form, by shifting cells to serum starvation, and subsequently induced to retract, by re-feeding serum. We stimulated (STI) WT & KO C2C12 cells to re-enter the cell cycle after serum starvation and quantitated the number of cells with cilia as well as their ciliary length. WT cells retract their cilia when refed, but KO cells were inefficient in resorption and continued to show large number of cells with cilia (Fig.5B & 5D). The length of cilia in serum fed KO cells was significantly longer than in WT cells. These results demonstrate that in addition to its role as an inhibitor of ciliogenesis, C3G functions to maintain cilia length, and aids in their resorption upon re-entry into the cell cycle.

### Catalytic activity of C3G is required for normal cilia formation

Since C3G has functions dependent on its catalytic activity as well as protein interaction, we wished to examine if the catalytic activity of C3G contributed to primary cilia maintenance. The presence of pC3G (the catalytically active form) at mitotic centrosomes suggested that the GEF function of C3G may be required at the centrosome. To address this, we used a dominant negative approach of expressing deletion constructs having only the central Crk binding region (CBR), or C3G lacking its catalytic domain ΔC-C3G (shown schematically in Fig.6A). Their expression was verified by western blotting (Fig.6B). These constructs have earlier been shown to inhibit catalytic activity dependent effector functions of cellular C3G (Guerrero et al., 1998; Martín-Encabo et al., 2007; Mitra and Radha, 2010; Radha et al., 2007; Radha et al., 2004; Sasi Kumar et al., 2015). CBR, as well as ΔC-C3G expressing cells show an increase in cilia formation (Fig.6C & 6D). CBR expressing cells also showed longer cilia similar to that seen in cells lacking C3G (Fig.6E). These results suggested that catalytic activity of C3G is required for maintaining primary cilia dynamics.

**Figure 6.**
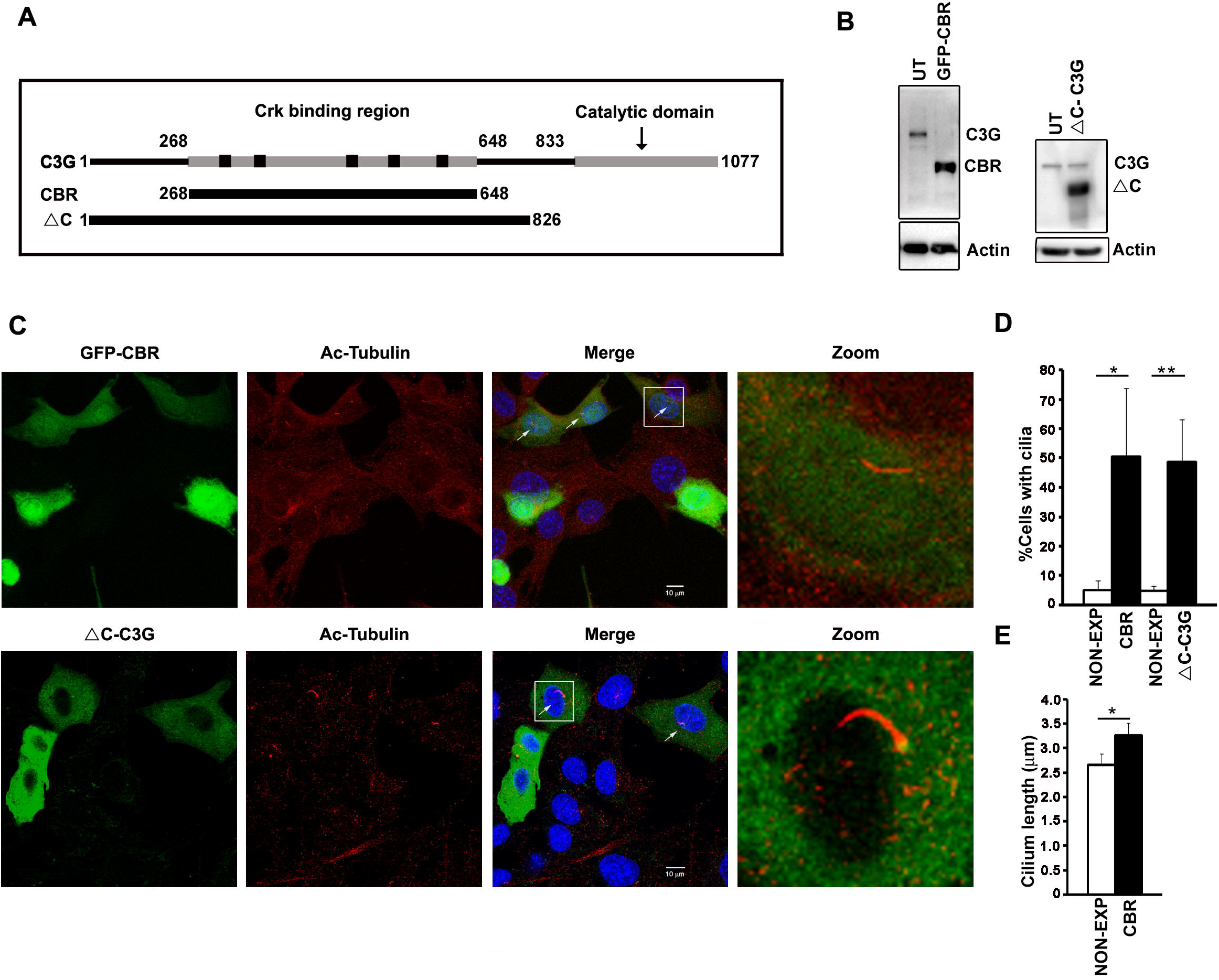
Inhibition of cellular C3G catalytic activity using dominant negative constructs enhances primary cilia formation. A. Schematic showing C3G deletion constructs lacking catalytic domain. B. C2C12 cells were transfected with GFP-CBR or ΔC-C3G, lysed and subjected to western blotting to show expression of the deletion constructs. Actin was used as a loading control. UT, untransfected cells. C. C2C12 cells transfected with GFP-CBR or ΔC-C3G were stained to detect expressing cells and Ac-tubulin. Arrows point to longer cilia seen in expressing cells. D. Bar graph showing proportion of expressing and non-expressing cells (NON-EXP) having cilia. *p<0.05, **p<0.01. E. Length of cilia in CBR expressing cells compared to that in non-expressing cells. *p<0.05.

### C3G is required for cell proliferation

Proteins that control centrosome division also regulate proliferation. While generating C3G KO clones, we observed that clones that have significantly low C3G expression grew slowly and did not survive beyond 2-3 passages. To examine if loss of C3G affects cell proliferation, we carried out assays to monitor growth of WT C2C12 and KO clones. Cell number was determined using MTT assays and proliferation was monitored over a period of 72 hrs. KO cells from the first passage proliferate very slowly compared to WT cells (Fig.7A). Cell cycle profile of exponentially growing cells from the first passage determined by FACS analysis showed that loss of C3G arrests cells in G1 phase (Fig.7B). KO cells from second passage showed increase in sub G1 population indicating that loss of C3G compromises cell survival (Fig.S6A). G1 arrest and delay in cell proliferation could be a consequence of changes in expression of genes that regulate cell cycle. We, therefore, compared the expression of cyclin A, D1 and E in WT & C3G KO cells and found that knock out cells have significantly lower levels of these three cyclins (Fig.7C). Reduction in cyclin D1 was also seen in a C3G KO clone of MDA-MB-231 cells (Fig.S6B).

**Figure 7.**
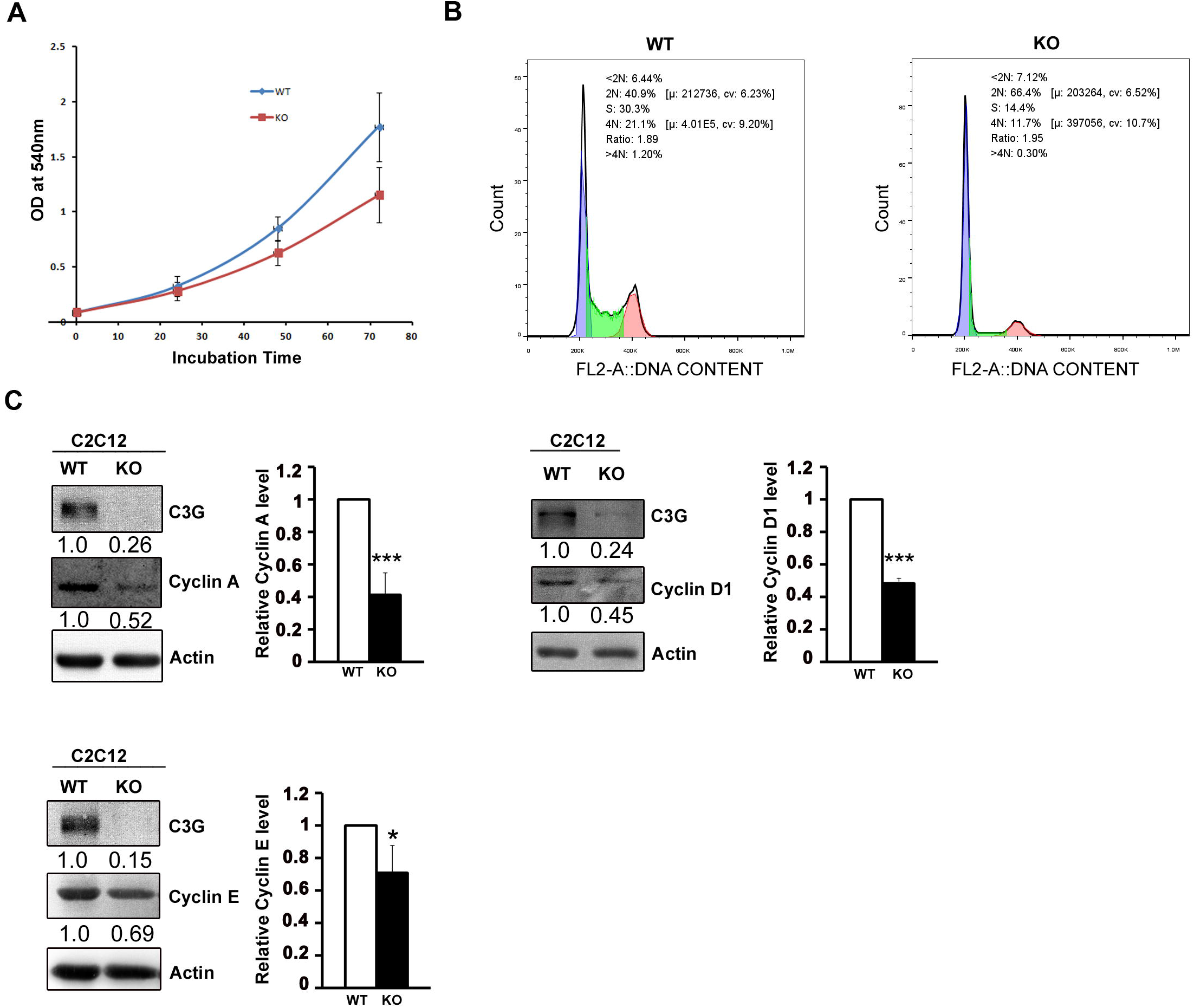
C3G Knockout cells show reduced Cyclin levels and proliferate slowly. A. Equal number of WT and KO C2C12 cells were seeded in a 96-well plate and subjected to MTT assay. Cell viability was assessed up to 72hrs at 24 hr intervals. Graph shows the absorbance measured at 540nm which reflects total number of live cells at each time point. B. Exponentially growing C2C12 WT and KO cells were stained with propidium iodide and analyzed by FACS for cell cycle profile. Figure shows representative flow cytometric profile of cells analyzed for DNA content indicating cells in various cell cycle phases. C. C2C12 WT and KO cell lysates were subjected to western blotting and probed for expression of C3G, Cyclin A, Cyclin D1 and Cyclin Actin was used as loading control. Respective bar graphs show the relative protein levels in KO cells compared to WT cells from three independent experiments. *p<0.05, ***p<0.001.

## Discussion

Through this study, we have identified C3G as a centrosomal protein and elucidated its role in regulating centrosomal division and primary cilia maintenance. The following evidences enabled us to conclude that C3G localizes to the centrosome and is required for maintaining centrosomal homeostasis as cells enter and exit the cell cycle: 1) C3G co-localizes with γ-tubulin and centrin in interphase as well as mitotic cells; 2) C3G interacts with and co-localizes with cenexin at the mother centriole; 3) C3G is present in the basal body of fully formed primary cilia; and 4) Cells lacking C3G due to CRISPR/CAS9 mediated knockdown show supernumerary centrioles and an increase in number of cells with cilia. KO cells do not show cilia retraction upon stimulation of quiescent cells to proliferate. In addition, we show that centrosomal localization of C3G is lost in differentiated myotubes, and that loss of C3G compromises proliferation with majority of the cells arresting in G1 and undergoing cell death.

The integrity of the centrosome, basal body and primary cilia are dependent on proper microtubule assembly. We found that C3G localization to the centriole is lost upon disruption of microtubules or microfilaments, suggesting that cytoplasmic C3G may be dynamically transported to the centrosome, aided by the cytoskeleton. Localization of other proteins, like geminin (which regulate centrosome duplication and ciliogenesis), to the centrosome is also dependent on an intact actin cytoskeleton (Lu et al., 2009). In addition, cilia length is controlled by regulators of the cytoskeleton (Copeland et al., 2018), indicating that microtubules and microfilaments can regulate cilia by localizing regulators to the centrosome, in addition to directly contributing to centrosome and cilia structure.

Many centrosomal proteins show cell cycle dependent localization to the centrosome (Hames et al., 2005; Kim et al., 2009; Lui et al., 2016). We observed that C3G particularly localizes to the mother centriole, with no detectable signal seen at the daughter centriole in interphase cells. In early prophase cells where the divided centrosomes have just split, C3G is seen associated with only one of the centrosomes, but late prophase cells show its presence in both. Sub distal appendage proteins are known to associate with the centrosome in late prophase (Guarguaglini et al., 2005; Nigg and Stearns, 2011; Vorobjev and Chentsov Yu, 1982). C3G locates to the newly divided centrosome around the time the mother centriole is marked by appendage proteins providing additional evidence for its association with appendages. We presently do not know why C3G shows increased centrosomal association during mitosis. Levels of proteins like PLK4 and SAS6, involved in building the PCM, are highest at the centrosome in mitotic phase and drop sharply at mitotic exit (Colicino and Hehnly, 2018). One possibility would be that increased association prevents ciliogenesis during mitosis. Negative regulators of ciliogenesis are required to suppress untimely formation of basal bodies in dividing cells (Kobayashi et al., 2011). Inhibition of ciliogenesis in dividing cells has been shown to be dependent on localization of proteins like Ndel1 to the sub distal appendages of the mother centriole (Inaba et al., 2016).

While C3G protein was associated with the centrosome in interphase cells as well as mitotic cells, tyrosine phosphorylated, or active form of C3G was seen at the centrosome only in mitotic cells. Also, its distribution was not restricted to the mother centriole, and appeared to be localized to the centrosome/PCM at large. pC3G was seen at the centrosome of interphase cells upon inhibition of tyrosine phosphatases, suggesting that C3G activation is very transient in interphase cells, and more stable at mitotic centrosomes. We have earlier shown that IMR-32 cells in interphase showed pC3G at the core of the Golgi upon stimulation by NGF, or Forskolin (Radha et al., 2008). It appears that C3G present at the centrosome may be specifically activated in a spatial and temporal manner at the onset of mitosis. Phosphorylation of proteins appears to be important for expansion of the PCM at the onset of mitosis (Guarguaglini et al., 2005; Haren et al., 2009; Ramani et al., 2018; Runkle et al., 2011). The targets of activated C3G at the centrosome require to be identified.

A very large number of proteins are associated with the centrosome, and a few show asymmetric localization to one of the centrioles (Alves-Cruzeiro et al., 2014; Jakobsen et al., 2011). Proteins localized to the mother centriole regulate centrosome integrity and division, as well as primary cilia dynamics, but not all of them have been identified, nor their functions well understood. Cellular C3G is seen predominantly in the cytoplasm, and undergoes regulated nuclear entry in response to physiological stimuli (Shakyawar et al., 2017). The localization of C3G to the centrosome and specifically to mother centriole is reported here for the first time. C3G specifically localized at one end of the γ-tubulin cylinder, perfectly co localizing with cenexin, a sub distal appendage protein. Since over-expression of C3G inhibits centriolar division and its loss results in formation of supernumerary centrioles, we concluded that C3G functions to maintain the centrosome division cycle and centriole numbers.

Cenexin regulates centrosomal cohesion by maintaining β-catenin levels, and restricts premature disjunction (Yang et al., 2018). C3G was earlier shown by us to function as negative regulator of β-catenin. Presence of C3G in a cellular complex with cenexin, and decrease in cellular cenexin levels seen in C3G depleted cells, suggest that C3G is required to maintain appropriate cenexin levels required to prevent supernumerary centriole formation. Whether C3G regulates cenexin expression at the transcriptional or post transcription level is to be examined. One possibility is that loss of C3G from the sub distal appendages of the mother centriole may trigger degradation of cenexin protein. Presently, very little information is available on regulation of cenexin protein levels.

The mother centriole forms the basal body which gives rise to the primary cilium. Molecular control of ciliogenesis has been investigated extensively, but our understanding of ciliary maintenance and resorption is still poor. Few molecules that localize to the mother centriole like CP110, CEP97, Ndel1 and APC, are negative regulators of ciliogenesis (Inaba et al., 2016; Spektor et al., 2007; Tsang et al., 2008; Wang et al., 2014). Also, molecules that bind tubulin and regulate centriolar/ciliary-microtubule construction are important determinants of cilia length (Zheng et al., 2016). We observed the presence of C3G at the basal body in cells that have put forth a primary cilium. The primary cilium is considered as a signaling hub enabling responses to chemical as well as mechano-sensory signals. C3G, is involved in signaling from growth factors as well as mechanical cues, and may therefore be participating in transducing signals received by the primary cilium (Marada et al., 2016; Radha et al., 2011; Takahashi et al., 2008; Tamada et al., 2004). In addition, our results implicate a function for C3G in maintaining the structure of the cilium, as loss of C3G resulted in longer cilia. Increased number of cells with primary cilia in C3G KO clones, or upon disruption of its catalytic activity by expression of dominant negative constructs is suggestive of its catalytic function in maintaining cilia length.

Retardation of ciliary resorption delays cell-cycle progression to the S and M phase after cell-cycle reentry (Kim and Tsiokas, 2011). Inability of cells lacking C3G to retract cilia upon stimulation to enter the cell cycle could be one of the reasons for slow rate of proliferation of the cells. We also observed that the clones could not be maintained beyond two passages. C3G knock out clones from the first passage arrest at G1 phase and significant cell death is seen in the second passage. We observed reduced levels of various cyclins required for G1 and S phase in knock out clones, that could also be responsible for their arrest at G1. We have earlier shown that C3G regulates chromatin modifications and gene expression (Shakyawar et al., 2017). C3G may, therefore, play a role in maintaining the balance between cell proliferation and quiescence by regulating gene expression as well as centrosomal division and primary cilia dynamics. Centrosomal localization of C3G is lost as myocytes fuse to form myotubes. Prominent nuclear localization of C3G is seen in myotubes. Our results, as well as work from other labs have shown that primary cilia are retracted, and centrosomes do not divide as myocytes fuse to form myotubes (Bugnard et al., 2005; Fu et al., 2014; Srsen et al., 2009). Since C3G regulates centrosomal division as well as primary cilium formation, irreversibly arrested cells may not require C3G function at the centrosome.

Several proteins deregulated in human cancers have been shown to affect centrosomal division and ciliogenesis (Bettencourt-Dias and Glover, 2007; Nigg and Holland, 2018; Plotnikova et al., 2008; Wang and Dynlacht, 2018; Gonczy, 2015). C3G has been implicated in tumorigenesis, with higher as well as lower levels seen in different cancers (Che et al., 2015; Guerrero et al., 1998; Gutiérrez-Berzal et al., 2006; Hirata et al., 2004; Okino et al., 2006; Priego et al., 2016; Radha et al., 2011; Samuelsson et al., 2011; Sequera et al., 2018). The function of C3G in regulating centrosome division may be responsible for its deregulation being associated with tumors.

Embryonic development is dependent on regulated division of each cell. Many molecules that are components of the centrosome or primary cilia are responsible for gross developmental disorders when mutated. Loss of C3G results in early embryonic lethality of mouse embryos, and this must be due to improper cell division and failure of timely differentiation. Regulation of centrosomal division and primary cilia dynamics are novel functions of C3G that provide molecular explanation for the early death seen in knock out embryos. We also hypothesize that mutations in C3G may cause specific developmental defects called ciliopathies in humans.

## Materials and Methods

### Cell culture, Transfections and Treatments

Mouse myoblast cell line C2C12, was cultured in Dulbecco’s modified eagle’s medium (DMEM), containing 20% fetal bovine serum. ARPE-19 cells were grown in F12-DMEM with 10% serum. IMR-32, HEK-293T & MDA-MB-231 cells were grown in DMEM containing 10% FBS, at 37°C and 5% C02. For myotube formation, C2C12 cells at 80% confluence were induced to differentiate by replacing FBS containing medium with media containing 2% horse serum for 72-96 hours (Kubo, 1991; Sasi Kumar et al., 2015). Lipofectamine Plus, LTX, 2000 and 3000 from Invitrogen were used for transfection as per manufacturer’s instructions. Treatments of cells were as follows: nocodazole (Calbiochem) 1μg/ml for 4 hours, Cytochalasin D (Calbiochem) 1μg/ml for 30 minutes, and hydroxyurea (Calbiochem), 3mM for 30 hours. For serum starvation (SS), cells were grown in media containing 0.5% serum for 24 hours. For restimulation (STI) to enter the cell cycle, starved cells were refed with 20% serum containing medium for 24 hours. C3G wild type (WT) and knockout (KO) C2C12 & MDA-MB-231 clones were generated using mouse and human specific CRISPR/CAS 9 constructs obtained from Santa Cruz as described earlier (Shakyawar et al., 2018). Cells from the first passage were used for all experiments unless otherwise mentioned. PV treatment was given as previously described (Radha et al., 2004).

### Antibodies & Plasmids

Antibodies were obtained from the following sources: Anti rabbit C3G (H300), sc-15359; anti mouse C3G (G4), sc-17840; anti rabbit pC3G (Tyr514, sc32621 & Tyr504, sc-12926); anti rabbit PCM1 (H262), sc-67204; anti rabbit Cyclin D1 (M-20), sc-718; anti rabbit Cyclin A (H-432), sc-751; anti rabbit Cyclin E (M-20), sc-481; anti mouse α-Tubulin (B-7), sc-5286 and anti mouse GFP (B2), sc-9996 from Santa Cruz biotechnology. Anti mouse Actin, MAB1501 was obtained from Millipore. Anti mouse γ-tubulin, T-6557 and anti mouse acetylated-tubulin, T-7451 were from Sigma-Aldrich. Anti rabbit Cenexin, ab43840 was from Abcam. Cells were stained for F-actin using Rhodamine phalloidin (Molecular Probes). Anti rabbit C3G-GRF2 (rGRF2), NBP1-88266 and anti mouse C3G-GRF2-2F5 (mGRF2), NBP2-45515 were from Novus Biologicals. C3G-GFP, C3G-Flag, GFP-CBR, ΔC-C3G, control (con-Sh) and ShC (ShRNA-C3G) plasmids have been described earlier (Dayma and Radha, 2011; Radha et al., 2007). GFP-Cenexin and GFP-Centrin were kindly gifted by Kyung Lee, NIH and Michel Bornens, Institute Curie, respectively. pEGFPC1, C3G CRSIPR/CAS9 KO Plasmids (mouse sc-430750 & human sc-401616), and C3G HDR Plasmid (mouse sc-430750-HDR & human sc-401616-HDR) were from Santa Cruz Biotechnology. Fluorophore conjugated secondary antibodies were from Millipore and Amersham GE. Adenoviral vector expressing human C3G was generated using the AdEasy System (Shakyawar et al., 2018).

### Immunofluorescence and Image Analysis

Cells grown on cover slips were fixed with cold methanol at −20°C for 6 minutes, washed with PBS and incubated with 2% BSA in PBS for 1 hour. Indirect immunofluorescence was carried out as described earlier (Shakyawar et al., 2017) and coverslips were mounted in mounting medium containing 4,6-diamino-2-phenylindole (DAPI). For dual labeling, cells were incubated with first primary antibody, then the corresponding secondary antibody, followed by the second primary antibody and finally, its corresponding secondary antibody. Unless otherwise mentioned, H300 antibody that detects C3G was used for all experiments. The processed cells were scanned on a Leica TCS Sp8 confocal microscope. Similar acquisition parameters were maintained during capture of images from samples belonging to the same experiment. Images were analysed for colocalization by measuring fluorescence intensity in an ROI drawn across the centrosome. Proportion of cells showing a primary cilium was quantitated by observing acetylated tubulin staining. Averages were obtained by examining multiple fields from each coverslip, of three-four independent experiments carried out in duplicate. Cilia length was determined from scanned images using the LAS X, version 3.1.1.15751 software embedded in the system provided by Leica. Cells expressing C3G, or its deletion constructs were examined for the presence of supernumerary centrioles by staining for γ-tubulin. Proportion of interphase cells with three or more γ-tubulin spots were considered as having supernumerary centrioles. Quantitative data was obtained by examining a minimum of 50 expressing, and 100 nonexpressing cells from each of the duplicate cover slips from 3 independent experiments.

### Immunoprecipitation & Western Blotting

Immunoprecipitation of over expressed GFP tagged protein was carried out using GFP trap (Chromotek) as indicated by the manufacturer. Immunoprecipitation of constructs lacking GFP tag was carried out using the corresponding primary antibody and Protein A/G plus agarose as described earlier (Mitra and Radha, 2010). Western blotting was carried out using standard protocols (Radha et al., 2007; Radha et al., 2004). Image J software was used for quantitation of band intensity and adjusted with loading control to estimate relative differences.

### MTT Assay

Equal number of wild type C2C12 and C3G knockout C2C12 cells were seeded in a 96 well plate in triplicates. Cell viability was assessed at 24-hour intervals for 72 hours. 20μl of MTT was added and incubated for 5 hours. Media was removed and the purple formazon crystals trapped in cells was dissolved by adding 100 μl of DMSO for 30 minutes and absorbance at 540 nm was measured using a plate reader.

### Cell Cycle Analysis

Cells were harvested as a single cell suspension in PBS and added slowly to a falcon tube containing 70% ethanol and incubated for 45 minutes. Cells were centrifuged and re-suspended in Propidium Iodide staining buffer (Ormerod, 1994), and processed for cell cycle analysis using Beckhman coulter Gallios flow cytometer. Cell cycle profiles were analyzed by FlowJo X software.

## Acknowledgements

We are grateful to Dr. Kyung Lee, NIH and Dr. Michel Bornens, Institute Curie, for gift of plasmids. We thank Dr. Jyotsna Dhawan, CCMB, for critical comments on the manuscript and Ms. Divya Sriram, CCMB for help with preparation of the manuscript and figures. We thank Mr. B.V.V. Parthasaradhi, CCMB, for help with MTT assay.

